# Early Emergence and Long-Term Persistence of HIV-Infected T Cell Clones in Children

**DOI:** 10.1101/2020.09.15.298406

**Authors:** Michael J. Bale, Mary Grace Katusiime, Daria Wells, Xiaolin Wu, Jonathan Spindler, Elias K. Halvas, Joshua C. Cyktor, Ann Wiegand, Wei Shao, Mark F. Cotton, Stephen H. Hughes, John W. Mellors, John M. Coffin, Gert U. Van Zyl, Mary F. Kearney

## Abstract

Little is known about the emergence and persistence of HIV-infected T cell clones in perinatally-infected children. We analyzed peripheral blood mononuclear cells for clonal expansion in 11 children who initiated antiretroviral therapy (ART) between 1.8-17.4 months of age and with viremia suppressed for 6-9 years. We obtained 8,662 HIV-1 integration sites from pre-ART and 1,861 sites on ART. Expanded clones of infected cells were detected pre-ART in 10/11 children. In 8 children, infected cell clones detected pre-ART persisted for 6-9 years on ART. A comparison of integration sites in the samples obtained on ART with healthy donor PBMC infected *ex-vivo* showed selection for cells with proviruses integrated in *BACH2* and *STAT5B*. Our analyses indicate that, despite marked differences in T cell composition and dynamics between children and adults, HIV-infected cell clones are established early in children, persist for up to 9 years on ART, and can be driven by proviral integration in proto-oncogenes.

## Introduction

Human Immunodeficiency Virus (HIV) remains a worldwide health crisis. Approximately 37.9 million people are living with HIV globally and about 1 million die each year (1). Although current antiretroviral therapy (ART) is able to fully suppress HIV-1 replication in the blood (2-5), lymph nodes (6-9), and other tissues (10, 11), it does not cure the infection. If treatment is initiated before the immune system is heavily compromised and if there is lifelong adherence, ART can lead to a partial restoration of CD4+ T cell numbers (12, 13) and can prevent immunodeficiency in most individuals.

The main obstacle to a cure for HIV-1 is the persistence of replication-competent proviruses in long-lived and/or proliferating populations of infected T cells (14, 15). Most of the infected cells that persist on ART contain defective proviruses that are incapable of producing infectious virus (16, 17), although they may be complemented to generate infectious virus upon ART interruption (18, 19). These defective proviruses do not directly contribute to the HIV-1 reservoir that persists on ART but complicate its measurement and may contribute to persistent immune activation. The fraction of infected cells that contains replication-competent (intact/infectious) proviruses has been estimated to be between 1 and 5% in individuals on long-term ART (16, 17, 20). Although the fraction of intact proviruses is small relative to the total number of infected cells, there are sufficient replication-competent proviruses, or defective proviruses that can be readily complemented, to fuel rapid viral rebound if ART is interrupted (21, 22). In both adults and children, when ART is initiated soon after infection, the number of infected cells is reduced, sometimes to levels below the detection limit of current assays (20, 23, 24) and rebound viremia can be significantly delayed (25-28).

Studies of HIV-1 integration sites were initially performed in cell lines and showed that sites were widely distributed but favored highly expressed genes (29-31). Two studies in 2014 were the first to demonstrate expansion of HIV-infected T cells *in vivo* (32, 33). These clones of infected T cells can be detected as early as Fiebig IV in acute infection (34), can persist in adults for at least 3 years on ART (35), and are distributed among different tissues (6). Studies of clones persisting in adults on ART revealed selection against proviruses in expressed genes with a stronger selection against those that are integrated in the same orientation as the host gene (32, 36) and selection for proviruses integrated into some proto-oncogenes—e.g. *BACH2, MKL2, and STAT5B* (32, 33). Although much is now known about the HIV-1 integration site landscape in adults prior to and on ART, there is little information on clonal expansion of infected cells in children who acquired HIV perinatally (PHIV). The largely anti-inflammatory and immunoregulatory environment of the immune systems in children (37, 38) could affect the behavior of infected T cells in ways that would alter the integration site landscape and selection of proviruses at specific sites in children, leading to differences compared to adults. Furthermore, infants have a high fraction of naïve T-cells and fewer clonally expanded T-cells than adults (39, 40), a difference that could affect integration site selection and clonal expansion of infected cells.

To our knowledge, only two reports have investigated clonal expansion of infected cells in children, to date (33, 41). However, only a few children were studied and the integration site sampling in these studies was shallow because it is difficult to collect large numbers of PBMC from infants and children. One study reported clones of infected cells in 3 children initiating ART during chronic infection (33) and the other in 3 neonates on ART who were followed for 2 years (41). Here, we expand on these studies to perform a deep look at the integration site landscape in 11 children initiating ART early and followed for 6-9 years of continual suppression of viremia. We performed a detailed analysis of the integration site landscape by comparing the findings to those in *ex vivo* infected adult PBMC and to those in infected adults on ART (6, 32). To study the emergence of infected CD4+ T cell clones before ART initiation, the dynamics of their long-term persistence, and their potential survival within select genes, we obtained 10,523 integration sites from CD4+ T cells in the perinatally-infected children using samples obtained prior to and during long-term ART. We compared longitudinal integration site datasets to look for evidence of long-term persistence of clones of infected T cells and to investigate the frequency and size of the infected cell clones in the children. Finally, to determine if there exists selective maintenance of infected cells within single genes, we analyzed the integration sites in children compared to sites obtained from *ex vivo-*infected, CD8-depleted PBMC (deposited at rid.ncifcrf.gov) (42, 43).

We report here that clones of infected cells are found in children as early as 1.8 months after birth and that some of the clones that arose early persisted for up to 9 years on ART. Strikingly, although there are noted differences between the immune environments in children compared to adults, our findings on the population of infected T cell clones are similar to what has been reported for adults, suggesting that clonal expansion is the main mechanism for persistence of HIV-1 in children whose viremia is suppressed by ART. We also found that the selection for proviruses integrated in certain genes is similar in adults and children and, importantly, that this selection occurs pre-ART. Integration events and selection for proviruses in these genes in children born with HIV-1 could have long-term effects in adulthood that have not been investigated and are not observed in adults who were not born with HIV infection.

## Results

### Participants and Sampling

PBMC were obtained from children enrolled in the Children with HIV and Early Antiretroviral therapy (CHER) randomized trial and post-CHER cohort (44) who were identified as plasma HIV-1 RNA positive by 7 weeks of age, initiated ART within 18 months of age (median: 5.1 months; range: [1.8 to 17.4] months), and had long-term, sustained suppression on ART (3) (Table S1). Children were included based on the availability of pre-ART PBMC and PBMC obtained after at least 6 years of continuous suppression of viremia (median: 8.1 years; range 6.8 to 9.1 years). The sex, pre-treatment plasma HIV-1 RNA, ART regimen, time to viral load suppression, and CD4 percentage after long-term ART are shown in Table S1. The pre-ART and on-ART enchriched-CD4+ T cells were analyzed for the presence and persistence of clones of infected cells. We obtained between 197 and 1386 (median: 655) integration sites from each of the samples taken before ART was initiated and between 77 and 432 (median: 137) integration sites from those after at least 6 years on ART (Table 1). In total, we obtained 10,523 HIV-1 integration sites from the 11 children.

**Table 1.** Number of Integration Sites and Infected Cell Clones Detected in Children Prior to and On ART.

| Donor ID | Age at ART initiation (months) | No. of integration sites obtained pre-ART<sup>a</sup> | No. of integration sites detected with >2 breakpoints pre-ART (>1 breakpoint) | Oligoclonality index<sup>b</sup> (pre-ART) | Years suppressed on-ART | HIV DNA cps/10<sup>6</sup> PBMC on-ART<sup>c</sup> | Number of integration sites obtained on-ART<sup>a</sup> | No. of integration sites detected with >1 breakpoint on-ART | Oligoclonality index<sup>b</sup> (on ART) | No. of Integration site matches between pre- and on- ART<sup>d</sup> |
| --- | --- | --- | --- | --- | --- | --- | --- | --- | --- | --- |
| ZA002 | 17.4 | 1064 | 27 (50) | 0.085 | 6.87 | 33 | 113 | 9 | 0.079 | 7 |
| ZA003 | 1.8 | 1386 | 7 (30) | 0.04 | 8.06 | 2 | 148 | 16 | 0.161 | 0 |
| ZA004 | 2.7 | 655 | 5 (13) | 0.024 | 7.92/8.76 | 24/-- | 255 | 25 | 0.223 | 3 |
| ZA005 | 6.0 | 583 | 2 (14) | 0.027 | 8.04 | 9 | 77 | 4 | 0.166 | 1 |
| ZA006 | 9.0 | 486 | 1 (6) | 0.012 | 7.45 | 47 | 137 | 20 | 0.313 | 1 |
| ZA007 | 9.9 | 197 | 4 (5) | 0.07 | 8.24 | 21 | 85 | 8 | 0.403 | 2 |
| ZA008 | 2.2 | 1293 | 1 (8) | 0.006 | 6.77 | 42 | 225 | 11 | 0.065 | 1 |
| ZA009 | 2.0 | 809 | 0 (16) | 0.021 | 9.13 | 186 | 125 | 5 | 0.055 | 0 |
| ZA010 | 1.8 | 514 | 1 (12) | 0.027 | 8.35 | 5 | 115 | 3 | 0.173 | 0 |
| ZA011 | 9.3 | 432 | 5 (9) | 0.037 | 7.35 | 182 | 149 | 8 | 0.092 | 3 |
| ZA012 | 5.1 | 1243 | 3 (12) | 0.01 | 8.41 | 12 | 432 | 32 | 0.126 | 2 |
| Median Values | 5.1 | 655 | 3 (12) | 0.027 | 8.04 | 24 | 137 | 9 | 0.161 | 1 |

<sup>a</sup>Value obtained by counting an integration from both the 5' and 3' LTR as a single integration site
<sup>b</sup>As described in Gillet, *et al.* and Bangham (41). This value ranges in the interval [0, 1] dependent on the relative size and contribution of integration site clones to the dataset where 0 signifies a completely uniform distribution while 1 signifies a single integration site.
<sup>c</sup>Integrase cell-associated DNA (iCAD) protocol (51)
<sup>d</sup>Matches between pre-ART and on-ART are counted as clones in columns 4 and 9.

### Clones of HIV-1 infected cells are detected in children pre-ART and persist on long-term ART

Clonal expansion of cells infected with replication-competent proviruses or defective proviruses that can be complemented during active replication (45, 46), is an important mechanism for HIV-1 persistence on ART (6, 14, 35, 47, 48). The detection of identical integration sites within a sample is the hallmark of clonal expansion of an infected cell, independent of the replication-competence of the integrated provirus. We defined an integration site as being from a clone using three separate criteria: 1) detection of the same integration site at least 3 times in pre-ART samples (to account for recently-infected cells that had duplicated their DNA but would die before establishing a clone), 2) detection of the same integration site at least twice in an on-ART sample (if a cell is dividing after long term ART, it is almost certainly part of a clone), and 3) detection of the same integration site in two different samples from the same donor. Additionally, the method we use to identify integration sites recovers the host-virus DNA junctions from both the 5’ and 3’ LTRs (32). Therefore, integration sites observed at both junctions were considered as a single integration site under the conservative assumption that they could have originated from the same provirus. In all but one of the 11 donors [Participant Identifier (PID) ZA009], we found at least one clone of infected cells in the pre-ART samples (range: [1 to 27]) (Table 1, column 4). We found at least 3 clones of infected cells in all on-ART samples (range: [3 to 32]) (Table 1, column 9). Although we did not detect any clones of infected cells in the pre-ART samples from donor ZA009 by the stringent criteria described above, 16 of the integration sites were detected twice, suggesting that clones of infected T cells could have been present in this donor pre-ART (Table 1, column 4 parenthetical). We identified clones of infected cells in the pre-ART samples that persisted for up to 6-9 years on ART in 8 of the 11 children (range: [1, 7] clones) (Table 1, column 11). These data show that clonal expansion contributes to the persistence of total HIV-1 DNA in children, as was shown previously for adults (6, 15, 32, 35, 47).

### Size of infected cell clones is similar in children and adults

We analyzed the size and frequency of infected cell clones using a modified Gini coefficient called the “oligoclonality index” (OCI) (49). Briefly, the OCI, which has a value between 0 and 1, is a measure of the non-uniformity of a given dataset; 0 indicates complete heterogeneity and 1 indicates complete homogeneity. In our analysis, 0 would mean each detected integration site was detected only once while a value of 1 would mean that all the integration sites would be from a single large clone. In the pre-ART samples, most integration sites were detected only once (Table 1, column 5). The pre-ART samples contained large numbers of recently infected cells that had not undergone clonal expansion. Thus, all pre-ART OCI values were less than 0.1 (range: 0.006 to 0.085; median: 0.027). The pre-ART OCI positively correlated with the age at which ART was initiated – presumably because clones increase in size with time, which makes it easier for us to detect them (Adj. R^2^=0.53; p=0.011) (Figure 1A). Stated differently, although clones can arise soon after infection (35, 50), they may require time to expand to a size that can be detected using the integration sites assay (35). As expected, the OCIs were significantly higher during long-term suppression on ART (range: 0.055 to 0.403; median: 0.161; p=0.002) (Table 1, column 10, Figure 1B), suggesting that the short survival of most recently-infected T cells makes it easier to detect clones of infected cells after long-term ART (32, 35). It should be noted; however, that the on-ART OCI does not correlate with time on ART (Adj. R^2^=-0.08; p=0.63), suggesting that clonal expansion during ART is not just a function of time, but rather a complex dependence on homeostatic, antigen-driven, and integration-driven proliferation (Figure 1B). We further compared the on-ART OCI in children to published datasets from 9 infected adults (6, 32) on long-term ART and found no statistical difference (p>0.99; Figure 2; numerical data found in Jupyter Notebook – see methods) (51).

**Figure 1.**
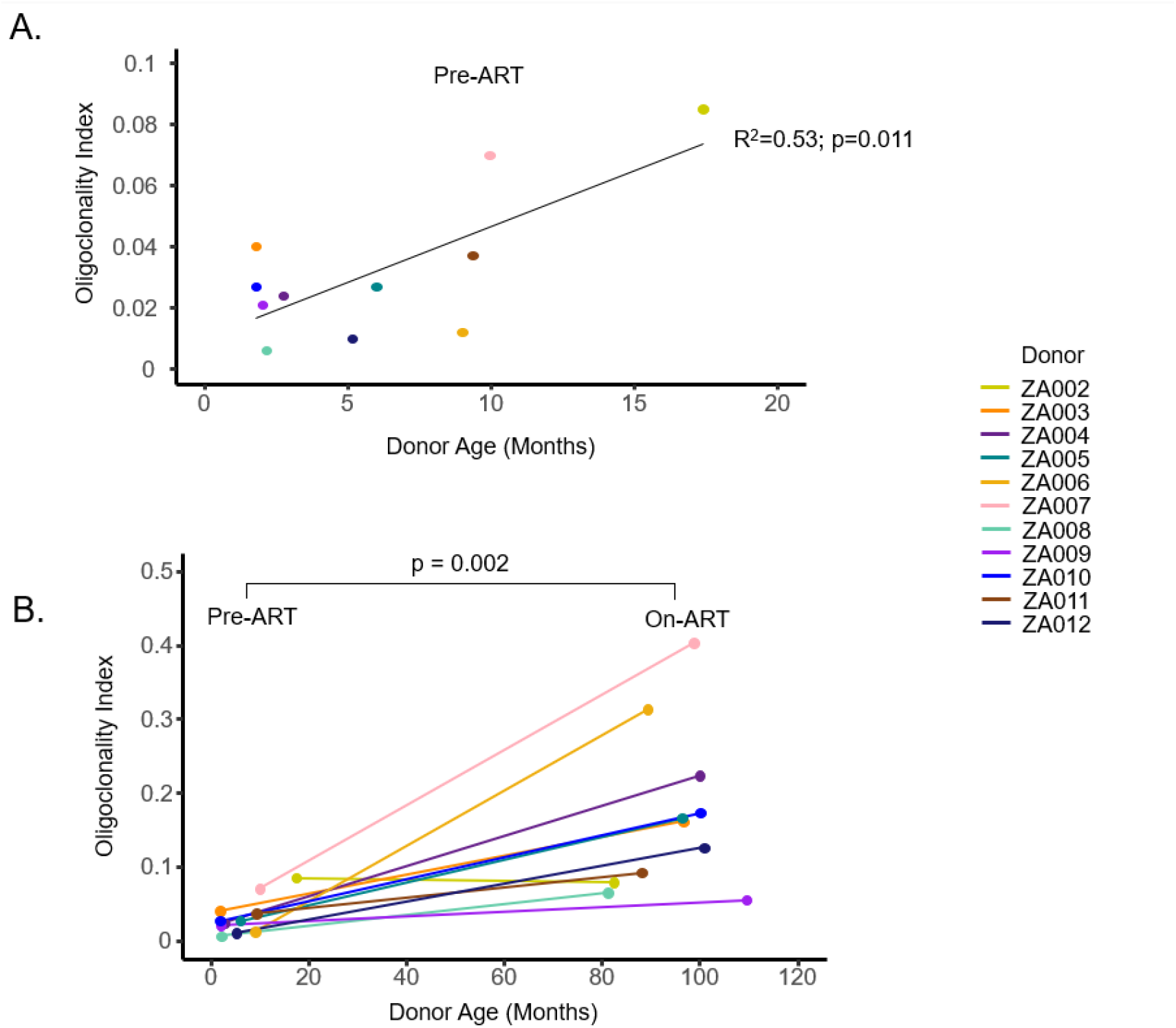
Oligoclonality indexes correlate both with time on ART and duration of infection prior to ART. Oligoclonality indexes (OCI) were calculated from the pre-ART and on-ART libraries plotted against Donor Age in months. Pre-ART OCIs were evaluated via linear regression and F-test against donor age **(A)** while change in OCI as a function of ART status was evaluated by Wilcoxon Signed-Rank test **(B)**.

**Figure 2.**
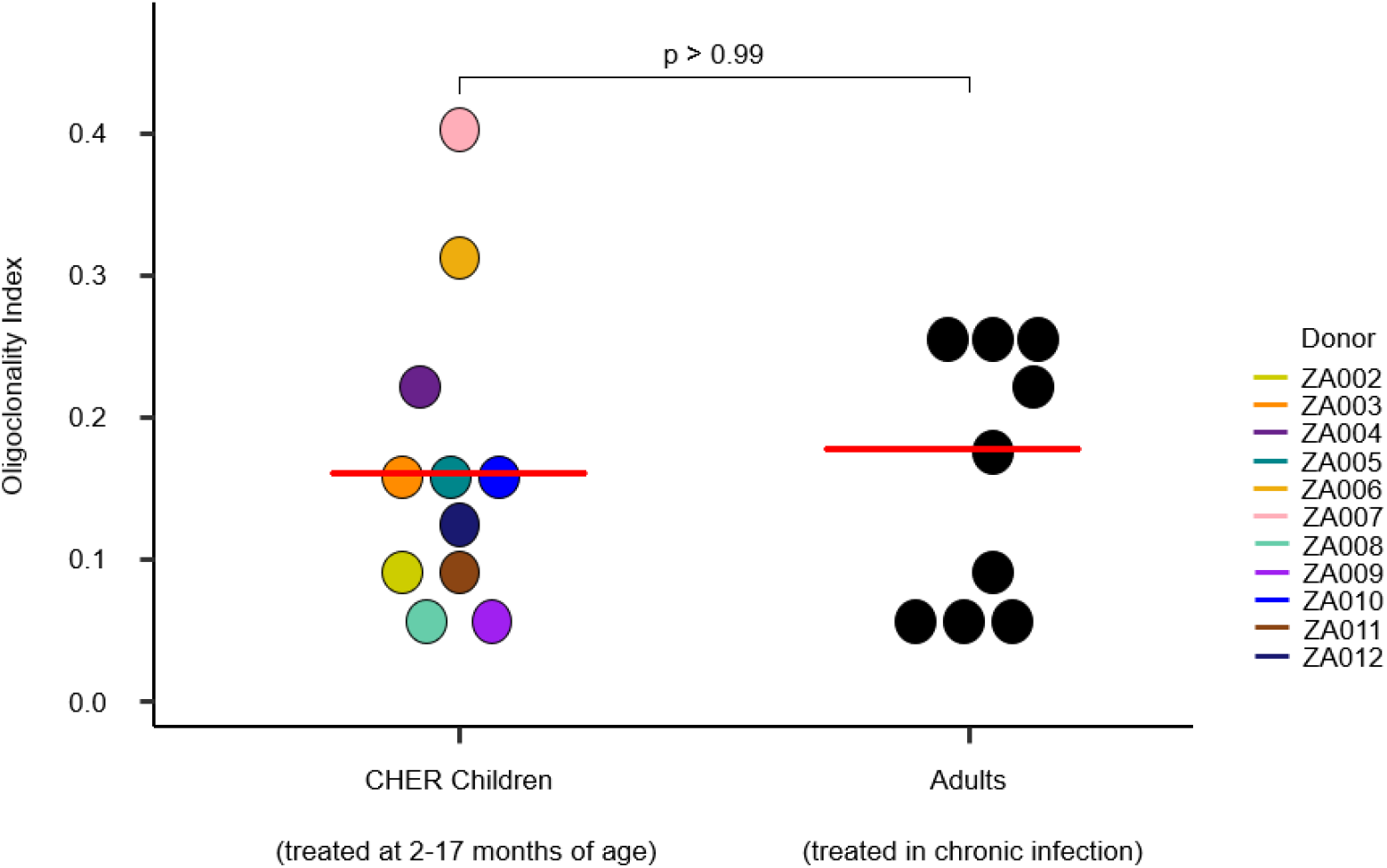
Oligoclonality indexes are comparable between ART-suppressed adults and children. Integration site data from donors whose viremia was suppressed on ART were downloaded from the Retrovirus Integration Database (rid.ncifcrf.gov) (43) from two studies totaling 9 individuals (6, 32) and the OCIs were calculated. OCIs were compared using Mann-Whitney test. Median values for each patient group are marked by red lines.

### Selection for cells with proviruses integrated in certain genes

Recent reports show that HIV-1 proviruses integrated in one of a small number of genes (15, 32, 33, 35, 52) contribute to the growth, survival, and persistence of the infected cell clones *in vivo*. To look for evidence of similar selection in children born with HIV-1 and treated early with ART, we compared the distribution of integration sites from the children (pre-ART and on-ART) to integration sites obtained from *ex vivo* HIV-1 infected, CD8-depleted PBMC from healthy donors [deposited at rid.ncifcrf.gov; (42)]. We asked if there was evidence for enrichment of proviruses in specific genes *in vivo* (relative to *ex vivo*). We also analyzed the orientation of the proviruses relative to the host gene. Enrichment in the fraction of proviruses within, and oriented in the same direction as, the gene are evidence of post-integration selection. Enrichment of the integration sites was determined by comparing the *ex vivo*-infected PBMC dataset against the *in vivo* datasets. For this analysis, clonally amplified sites were removed from the in vivo datasets by collapsing identical integration sites. Integration sites in intergenic regions (mapped to hg19) were not included in the analysis. The resulting datasets consisted of 335,614 integration sites from the *ex vivo* infected PBMC (87.2% of the initial data), 7039 sites from the pre-ART dataset from the children (83.9%), and 1202 (76.8%) sites from the on-ART dataset from the children. To detect enrichment in both the pre-ART and on-ART datasets relative to the *ex vivo* PBMC dataset, Fisher’s Exact Tests were performed on genes in each library with post-hoc multiple tests correction. Adjusted p-values are reported with p_adj_ ≤ 0.05 being considered significant.

Consistent with what has been observed in virally suppressed adults (32), we found a strong enrichment for proviruses during ART integrated into both *BACH2* (p_adj_=2.7*10^−15^) and *STAT5B* (p_adj_=4.0*10^−29^) (Table 2), but not *MKL2* (p_adj_>0.05) during ART. The question of enrichment in samples prior to ART initiation in either adults or children has not previously been addressed. Strikingly, we observed a signal for enrichment of integrations into *BACH2* in children even prior to ART initiation (p_adj_=8.9*10^−17^) showing that selection can occur early in PHIV infection. Although not statistically significant, we also observed a trend toward selection for integration events in *STAT5B* (p_adj_=0.14) (Table 2) prior to ART initiation.

**Table 2.** Analysis of Enrichment of Integration into Specific Genes in vivo^a^

| Chromosome | Gene name<sup>b</sup> | Independent integrations *ex vivo*<sup>c</sup> | Independent integrations in CHER cohort pre-ART | Adjusted p-value<sup>d</sup> | Independent integrations in CHER cohort on-ART | Adjusted p-value<sup>d</sup> |
| --- | --- | --- | --- | --- | --- | --- |
| 17 | *STAT5B* | 562 | 29 | 0.14 | 37 | 4.0E-29 |
| 6 | *BACH2* | 132 | 31 | 8.9E-17 | 16 | 2.7E-15 |
| All | All other genes | 334,920 | 6,979 | >0.05 | 1,149 | >0.05 |

<sup>a</sup>Data shown only for integrations into genes and for which at least 1 integration was detected in both libraries
<sup>b</sup>Genic coordinates mapped to hg19
<sup>c</sup>*Ex-vivo* dataset contains integration sites from CD8-depleted PBMCs from two healthy donor patients infected and PHA-stimulated *ex vivo*
<sup>d</sup>Adjusted p-value determined by Fisher's Exact Test with post-hoc Benjamini-Hochberg Correction

Previous studies in adults have shown that, if there is post-integration selection for an HIV provirus in a gene, like STAT5B and *BACH2*, the proviruses are highly enriched for the same orientation as the gene (32, 33). We analyzed the genes for which there were at least 15 unique integrations in the *ex vivo* dataset and at least 1 integration in the *in vivo* dataset so that there would be a signal sufficient to detect selection. Although 18 genes were retained for analysis in the pre-ART dataset, only 2 met these criteria in the on-ART dataset (Table S2, S3). Despite the global preference for proviruses detected on ART to be integrated against the gene (*ex vivo* PBMC: 50.0% vs. children on-ART: 54.7%; p=0.0011), there was no evidence for such global selection prior to initiation of ART (*ex vivo* PBMC: 50.0%; children pre-ART: 50.7%; p=0.26 for the difference) (Figure 3A). However, of the 18 genes in which there were sufficient numbers of integrations in pre-ART samples, we found selection for with-the-gene integration in both *BACH2* (p_adj_=2.0*10^−3^) and *STAT5B* (p_adj_=7.8*10^−3^) (Table S2, Figure 3B) and an against-the-gene bias in an ankyrin repeat protein, *ANKRD11* (p_adj_=0.028) (Table S3). Although these data provide evidence for strong selection for both *BACH2* and *STAT5B* pre-ART, we do not consider the against-gene bias for *ANKRD11* to be evidence of selection specific to that gene because of the global bias for against-gene integrations and the lack of an enrichment signal in this and previous datasets.

**Figure 3.**
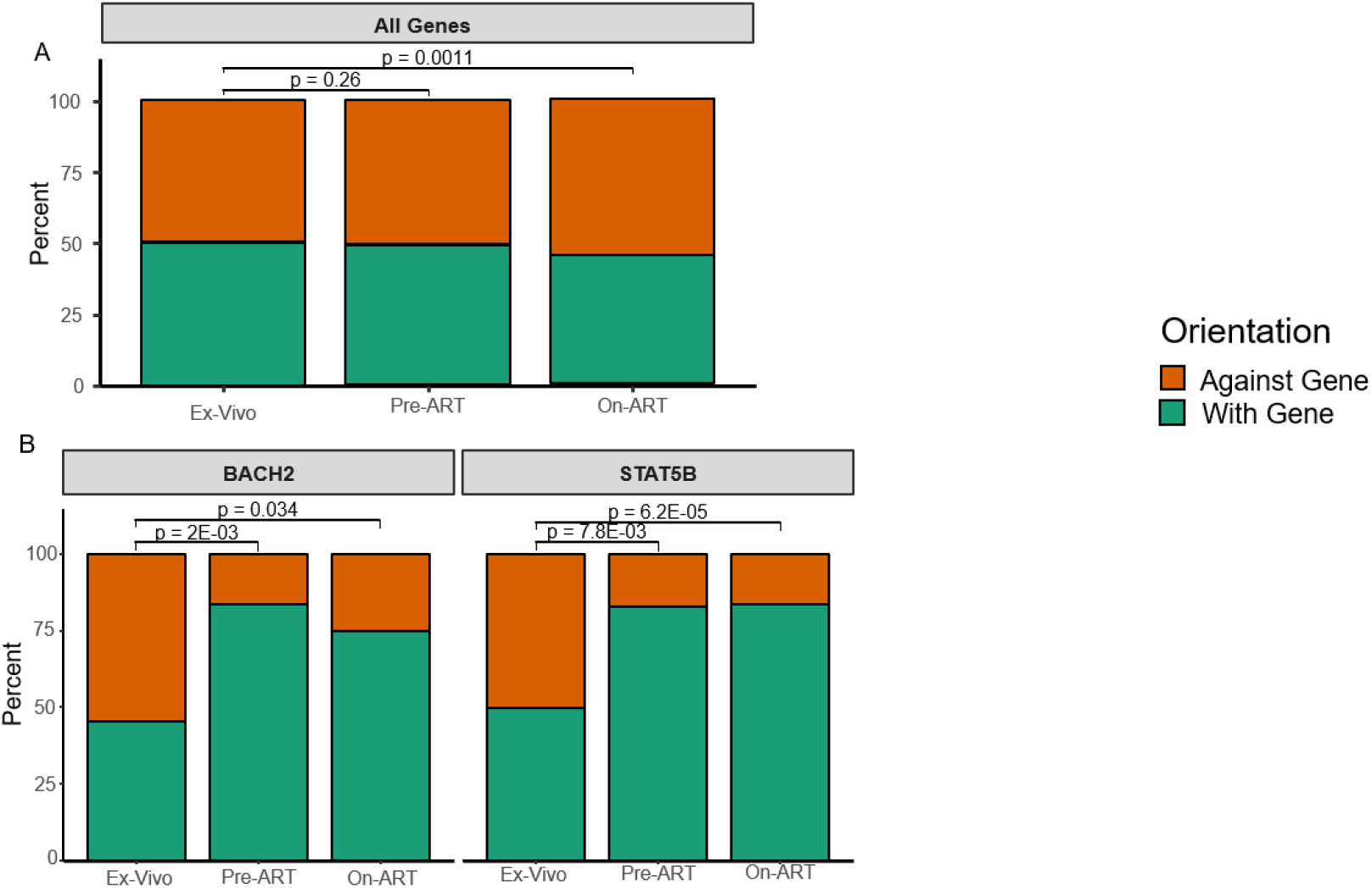
Global selection against proviruses oriented with-the-gene and selection in two genes for with-the-gene proviruses. For each of the integration site libraries, unique integration sites for all genes **(A)** and for proviruses integrated in *BACH2* and *STAT5B* **(B)** were plotted as the percentage of integrations against-the-gene (orange) and with-the-gene (green). Significance was assessed via Fisher’s Exact test between the *ex vivo* infected PBMC library and the pre-ART and on-ART integration site libraries from children. p-Values for pre-ART comparisons were post-hoc adjusted. The on-ART comparisons were not adjusted because of the differences in the number of independent statistical tests against each library.

Likewise, integration sites recovered from children on ART in *BACH2* and *STAT5B* were significantly selected for with-the-gene orientation (p_*BACH2*_=0.034; p_*STAT5B*_=6.2*10^−5^) (Table S3, Figure 3B). Taken together with the enrichment analyses, we conclude that cells containing proviruses integrated in *BACH2* and *STAT5B* in the same orientation as the genes were selected in children both prior to and on-ART.

We also compared the within-gene distribution of proviruses in *BACH2* and *STAT5B* in the children vs. the *ex vivo*-infected PBMC using an in-house mapping application (36) (Figure 4). In both genes, clearly visible clusters of integration sites in the same orientation as the gene (shown in blue) in a single intron upstream of the start of translation were observed in the children both before and during ART (Figure 4B, C, E, F). The *ex vivo*-infected PBMC have a broader, randomly oriented (equal red to blue) distribution (Figure 4A, D) in comparison. The different distributions highlight the selection for directional and clustered integration events into *BACH2* and *STAT5B* in children both prior to and on-ART.

**Figure 4.**
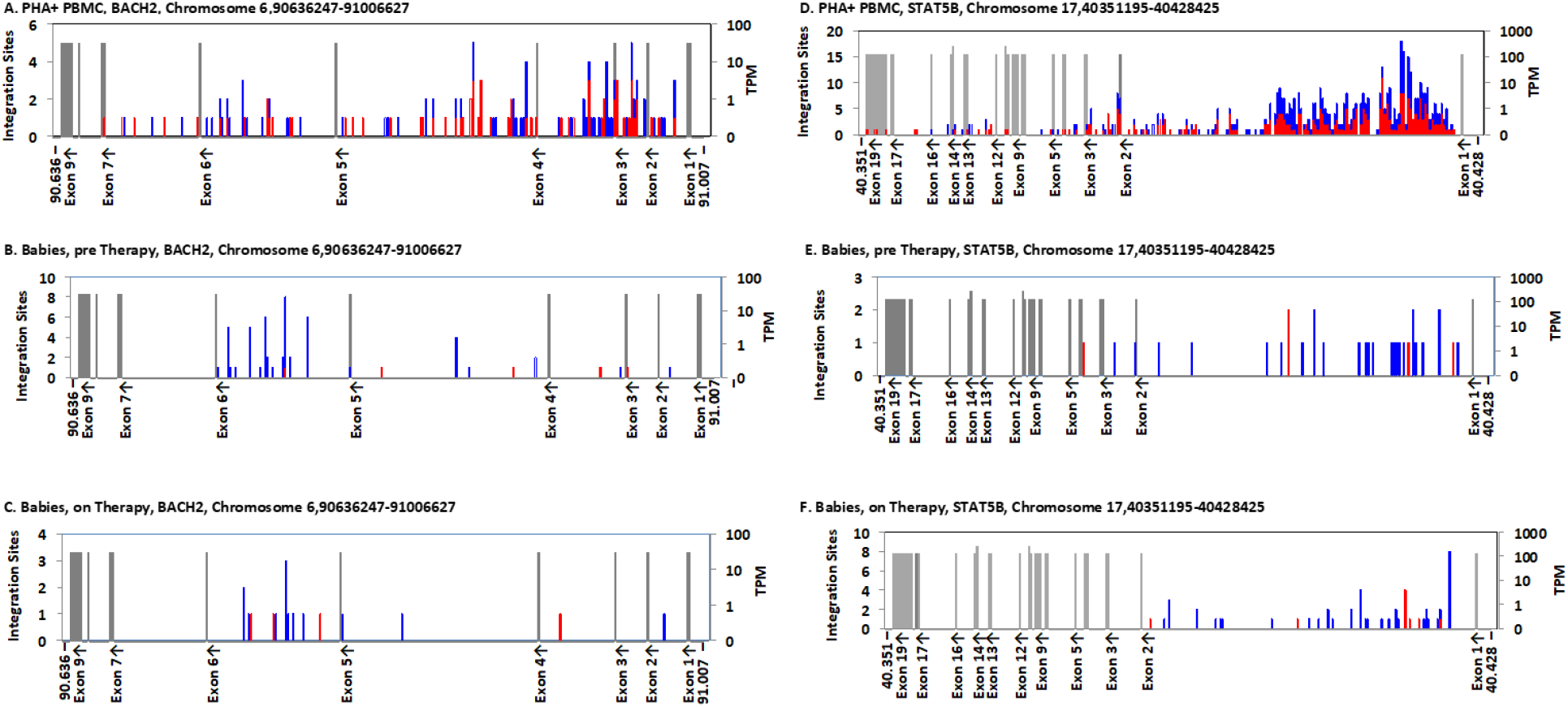
Distribution of Integration sites in *BACH2* and *STAT5B*. The maps show the integration sites of the proviruses in *ex vivo* infected PBMC **(A, D)** (36), infants sampled pre-ART **(B, E)**, and children sampled on ART **(C, F)**. Each graph shows the entire gene, divided into 250 bins. For *BACH2* **(A-C)**, each bin corresponds to ca 1500 NT; and for *STAT5B* **(D-F)**, ca 300 NT. Exons (labeled on the X axis, with orientation of transcription shown) are shown as grey bars, whose height indicates the level of expression, in transcripts per million (TPM), as shown on the scale on the right. Note that the resolution of the text sometimes leads to loss of labels of closely spaced exons. The numbers of integration sites in each bin are indicated by the stacked bars, according to the scale on the left, with red indicating the same transcriptional orientation as the chromosome numbering and blue indicating the opposite orientation. In these two genes, blue indicates the number of proviruses in each bin integrated in the same orientation as the gene.

### Sub-genomic sequencing datasets do not accurately characterize clonality within individuals

Proviruses in a subset of the children in this study were previously characterized using single-genome sequencing (SGS) of the *gag-pol* genes (encoding P6, protease, and the first 900 nucleotides of reverse transcriptase) (3). We assessed the clonality of the infected cells using integration site analysis compared to the identical sequences found in the SGS analysis. We found that proviruses with identical sub-genomic sequences were more common and constituted larger fractions of the data than the clones detected by sequencing integration site analyses (Figure 5, Figure S1). We also calculated the OCI for each set of data and found that the OCIs were significantly higher (average fold difference: 3.4x) for the sub-genomic single-genome sequences than for the integration site datasets (p = 0.0078) (Figure 5). These data suggest that either proviruses with identical sub-genomic sequences have different sites of integration, as has been shown for adults (47), or that many of the integration sites that were detected contained proviruses for which the *gag-pol* regions could not be amplified and sequenced due to deletions, PCR primer mismatches, or both.

**Figure 5.**
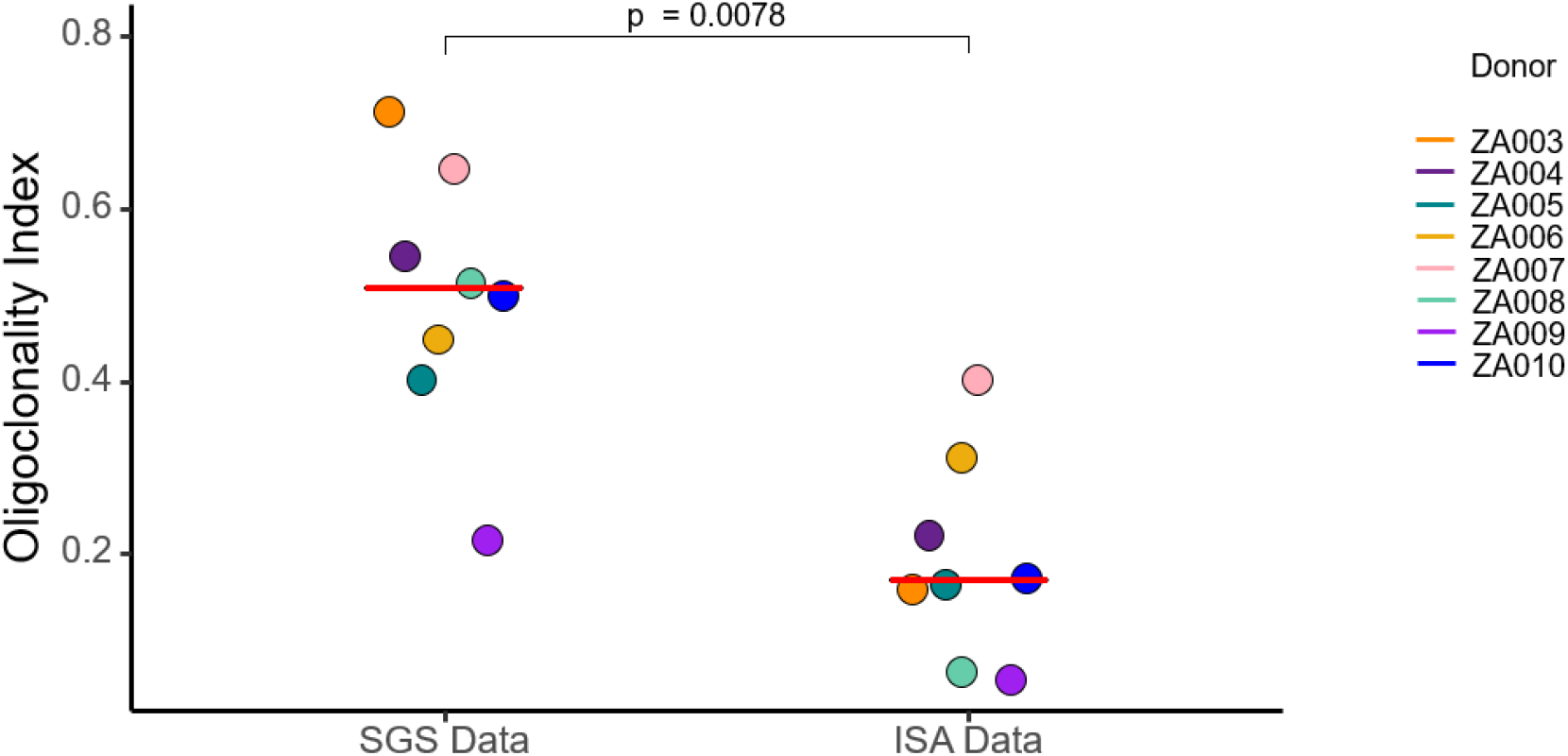
OCIs for single-genome sequencing datasets are significantly higher than OCIs derived from integration sites analyses. OCIs were calculated from single-genome sequencing and integration sites data obtained from PBMC of children suppressed for 6-9 years on ART. Significance was assessed by Wilcoxon Signed-Rank Test. Median values are noted by red dash for each group.

## Discussion

Despite effective therapies, which have reduced the rate of mother-to-child HIV-1 transmission (53-55), approximately 180,000 infants were infected worldwide in 2018 (53). These children must be included in the larger quest for effective HIV-1 curative interventions and such interventions may need to be tailored to their developing immune systems. Although the contribution of clonal expansion to HIV-1 persistence is well-studied in adults (6, 32, 35), this mechanism has not been well-described in children. Additionally, no analysis has been done in children on the clonal expansion of infected cells prior to the initiation of ART. To compare the mechanisms that underlie the persistence of HIV-1 infected cells during ART in adults and vertically-infected children, we performed HIV-1 integration site analysis on samples obtained from perinatally infected infants (prior to ART initiation) and from the same children during long-term suppression of viremia on ART (6-9 years of full suppression on ART). Despite inherent differences in T cell composition between children and adults (40) the clones of HIV-1 infected cells obtained from the blood of PHIV children in our study were not statistically different from adults (6, 32).

A study by Coffin et al. showed that infected cell clones can arise in adults in the first few weeks post-infection (35). In this study, we found that infected cell clones were detectable, using the integration sites assay, in 4 of the 5 samples collected from infants <3 months of age, consistent with early detection of clones in adults (35). In 2 of the 5 infants first sampled at <3 months old, we detected multiple proviruses with identical integration sites in both the pre-ART sample and the 6-9 years on-ART sample, demonstrating that clones of cells arose prior to ART initiation and persisted for years on ART. The other 7 donors, who were >3 months of age when initiating ART, also had detectable infected cell clones that persisted for at least 6 years of treatment. The frequency of clonal detection in the pre-ART populations tracked linearly with the estimated duration of infection prior to ART – using age as a surrogate – suggesting that the number of infected cell clones that expanded to detectable levels increased with the time of untreated infection, at least during the relatively short periods our donors were infected pre-ART. Our finding that infected cell clones had expanded and become large enough to be detected before two months of age supports the idea that the HIV-1 reservoir is generated rapidly, in actively dividing cells, in both adults and children (35, 56).

These results, in conjunction with previous studies showing that ongoing HIV-1 replication does not occur in children when viremia is fully suppressed on ART (3, 57, 58) and the fact that intact proviruses persist for years both in adults treated early (16) and in children treated early (20), supports the conclusion that the HIV-1 reservoir is maintained in vertically-infected children through the proliferation of cells infected prior to ART initiation, as it is in adults (6, 32, 35, 59). However, the available data are limited by the rarity of infected cells and the very small subset of HIV-infected cells that harbor intact, replication-competent proviruses in children (20). Although further studies are required to increase our understanding of the clonal expansion of intact proviruses as a mechanism by which the reservoir persists in both children and adults, it is possible that defective proviruses can undergo complementation upon ART interruption and contribute to viral rebound (60).

Although the number of infected cells in children on ART is small, we were able to detect an enrichment in the number and the orientation of proviruses in both *BACH2* and *STAT5B* in the pre-ART and on-ART samples, suggesting that proviruses in a specific intron and oriented with these genes can promote the survival of these clones *in vivo*, as in adults (32). While the selection for the survival of cells harboring *BACH2* and *STAT5B* proviruses has been previously described in adults on-ART (32, 33), no data had been presented to show that such selection exists prior to ART initiation. In both pre- and on-ART, we saw clear evidence for selection of cells containing proviruses in the exon immediately upstream of the start site of translation in *BACH2* and *STAT5B*. Although the selection of BACH2 integrants pre-ART was largely driven by a single child (ZA002) who did not initiate ART until 17 months of age, this single example nonetheless shows that clonal selection due to integration in specific genes is not strictly an on-ART phenomenon. The duration of untreated infection in this child may have allowed enough time for the selection of the cells with the *BACH2* proviruses to become detectable. Similar conclusions can be drawn for selection for proviruses integrated in the first intron of STAT5B, where there was clear evidence of selection for cells containing proviruses in the first intron, despite it’s being a very strong target for integration *ex vivo*. The trend towards enrichment of *STAT5B* integrants in pre-ART samples was due to the high level of sampling required to overcome the background of integration events in this gene compared to the *ex vivo*-PBMC infected dataset; however, the statistically significant orientation bias prior to ART demonstrates that pre-ART selection exists for *STAT5B*.

Samples from a subset of the children studied here were previously characterized in experiments that showed that ART is effective in suppressing on-going cycles of viral replication in children (3). Thus, proviral SGS data were available at the same on-ART timepoint. The OCIs obtained using the P6-PR-RT SGS results were significantly higher than the OCIs obtained from the on-ART integration site data. The observation that a higher OCI was obtained from the SGS data than the ISA data adds to the growing number of studies (15, 47, 50) suggesting that viruses with identical sub-genomic sequences may not all come from a clonal population of infected cells. These data strongly suggest that sub-genomic sequencing does not always accurately identify clones of infected cells or sufficiently characterize the genetic diversity of the intra-patient HIV-1 populations that persist on ART (47). Although the results here are consistent with previous studies showing that sub-genomic sequences are not sufficient to define clonality, it should be noted that calculating an OCI for small-N datasets can result in artificially high OCI values. Studies that are based on integration site analysis, rather than SGS, are more appropriate to study the clonal expansion of infected cells.

It is important to note that because these children were diagnosed within a few weeks of birth it is not known whether the transmission of HIV-1 occurred at birth or in utero. Because of this ambiguity, the age of the participant may not accurately reflect the duration of infection, although we found evidence of clonal expansion as early as 1.8 months after birth. Furthermore, the integration site libraries only represent a small fraction of the total number of infected cells in the blood. It is therefore likely that many of the integration sites that were recovered only once belong to clones of infected cells.

Despite these caveats, we have presented here the largest dataset yet of integration sites from pediatric HIV-1 infections both prior to ART and after durable suppression on ART. Because children primarily have naïve T cells, which do not have the HIV coreceptor CCR5 as a surface marker (40), as well as an immune environment that promotes quiescence (37, 38), and a more diverse T cell receptor repertoire (39), it is important to determine if there are differences between the observed frequency of clones and patterns of integration and post-insertional selection in children and adults. However, despite the differences in the immune systems of adults and children, our data suggest that these differences do not influence the infection and clonal expansion of T cells to a degree that is detectable by our integration site analysis. It is possible that by 6 to 9 years of age the immune system may be similar enough to that of an adult to account for the striking similarities in the on-ART libraries of these children and the published data from adults. Although these data suggest that the role of clonal expansion as the mechanism for HIV-1 persistence during ART is similar in children and adults, further studies are warranted to better understand how the developing immune system affects clonal expansion and what effects proposed curative interventions might have in both children and adults.

## Materials and Methods

### Study Approval and Ethics Statement

The CHER trial is registered with ClinicalTrials.gov (NCT00102960). Guardians of all donors provided written informed consent and the study was approved by the Stellenbosch University Internal Review Board.

### Total HIV-1 DNA quantification

HIV-1 DNA levels were determined using the integrase cell-associated DNA (iCAD) assay as previously described (61) with the following primers for use with HIV-1 Subtype C:

Forward primer HIV_Int_FP CCCTACAATCCCCAAAGTCA 4653 → 4672

Reverse primer HIV_Int_RP CACAATCATCACCTGCCATC 5051 → 5070

### Integration Sites Assay

ISA was performed and analyzed as previously described (32, 62) using patient-specific primers to the 5’ and 3’ LTRs. Importantly, our protocol includes a shearing step (63) that effectively tags each DNA molecule, allowing determination of the relative numbers of cells in the initial pool with identical sites of integration (i.e., clonality). The full set of integration sites obtained has been submitted to the Retroviral Integration Sites Database (https://rid.ncifcrf.gov/) (43) and the primer sequences are available in Supplemental Table 5.

A comparison integration site dataset was prepared from CD8-depleted PBMC isolated from two HIV negative human donors infected *in vitro* with replication-competent HIV-1, subtype B (BAL) (64). After 2 days the cells were harvested and DNA was prepared and integration sites analyzed as previously described (36). The global distribution of the integration sites from the two donors, was indistinguishable; therefore all comparisons were performed with combined data from the two donors.

### Oligoclonality Index

The oligoclonality Index (OCI) was calculated using a python script available at https://github.com/michaelbale/python_stuff/. Full details of the calculation are described in the supplemental text of Gillet, *et al* (49). Briefly, the LTR-corrected counts of all unique integration sites are sorted into descending order and the cumulative abundance of the clones are summed as a fraction of the total number of unique integration sites and normalized to have a maximal value of 1. Mathematically, the OCI is calculated as below:

*s*_*i*_ – LTR-corrected count of integration site *i*

*S*– Number of unique integration sites in library

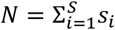– Total number of integration sites in library

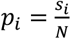– Relative abundance of integration site *i*

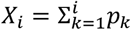– cumulative abundance of all integration sites of size {*s*_*i*_}or greater

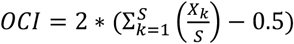– Olignoclonality index of library

### Statistical Analysis

Clonality was assessed by grouping sequenced integration sites with identical pseudo-3’LTR genomic coordinates and different shear points into count data in R. These count data were used to generate the OCI. Independent integration sites into genes were pooled and assessed for selection by Fisher’s Exact test by either the pre-ART or on-ART library vs. the ex vivo infected PBMC library as null set. P-values for this gene-enrichment analysis were corrected post-hoc by the Benjamini-Hochberg method. Orientation biases were assessed in a similar manner with post-hoc corrections only in the pre-ART comparison. All adjusted p-values are presented as p_adj_ where appropriate. All other statistical analyses are noted where appropriate and performed in R v3.5.2. A Jupyter notebook (51) with the R commands and visualizations for the unedited figures available at github.com/michaelbale/cher_bale/.

### Phylogenetic Analyses

HIV-1 P6-PR-RT sequences were aligned to HIV Consensus C using MUSCLE and neighbor joining phylogenetic p-distance trees were built using MEGA 7 (https://www.megasoftware.net/) (65) and outgroup rooted to Consensus C. Distance matrix generation for calculation of the sequence-based OCI was performed using Hamming distance.

## General

We thank the participants and their caregivers for their willingness to take part in this study as well as the clinic staff for their assistance.

## Author Contributions

MJB – Analyzed data, performed statistical analyses, wrote the paper

MGK – Processed samples, performed SGS, analyzed data, wrote the paper

DW – Performed ISA, analyzed data

XW – Performed ISA, analyzed data

JS – Processed samples, performed SGS

EKH – Processed samples, performed DNA quantitation

JCC – Performed DNA quantitation

AW – Performed SGS

WS – Analyzed Data

MFC – Conceived of idea, reviewed manuscript

SHH – Conceived of idea, analyzed data, wrote the paper

JWM – Conceived of idea, wrote the paper

JMC–Analyzed data, wrote the paper

GUVZ – Conceived of idea, wrote the paper

MFK – Conceived of idea, analyzed data, wrote the paper

## Data and materials availability

Previously published sequences from Van Zyl et al (3) were accessed from GenBank with accession numbers (KY820119-KY820376) retaining only the sequences associated with the on-ART timepoint of PIDs ZA003 – ZA010. The integration site libraries for the donors in this study are available in the Retrovirus Integration Database (RID) (43) at rid.ncifcrf.gov.The ex vivo infected PBMC integration sites library is also available in the RID under the Pubmed ID 31291371 (deposited at rid.ncifcrf.gov) (42).

## Supplementary Materials

**Table S1.** Donor Characteristics.

| PID | Sex | Time to viral load suppression in years | Pre-ART plasma HIV RNA<sup>a</sup> | ART regimen<sup>b</sup> | Years suppressed on ART | CD4% at on-ART time point |
| --- | --- | --- | --- | --- | --- | --- |
| ZA002 | Male | 1.37 | 654000 | AZT/3TC/LPV/r | 6.87 | 33 |
| ZA003 | Male | 0.46 | >750000 | ABC/3TC/LPV/r | 8.06 | 21 |
| ZA004 | Male | 1.38 | >750000 | AZT/3TC/LPV/r | 7.92/8.76 | 46 |
| ZA005 | Male | 0.47 | >750000 | AZT/3TC/LPV/r | 8.04 | 41 |
| ZA006 | Female | 0.44 | 635000 | AZT/3TC/LPV/r | 7.45 | 50 |
| ZA007 | Male | 0.92 | >750000 | AZT/3TC/EFV | 8.24 | 35 |
| ZA008 | Female | 0.44 | >750000 | AZT/3TC/LPV/r | 6.77 | 36 |
| ZA009 | Female | 3.76 | >750000 | AZT/3TC/EFV | 9.13 | 29 |
| ZA010 | Female | 0.46 | 510000 | AZT/3TC/LPV/r | 8.35 | 39 |
| ZA011 | Female | 2.29 | >750000 | AZT/3TC/LPV/r | 7.35 | 54 |
| ZA012 | Male | 0.93 | 277,000 | AZT/3TC/LPV/r | 8.41 | 31 |

<sup>a</sup>Determined by Roche Amplicor HIV Monitor assay v1.0
<sup>b</sup>ART abbreviations: Zidovudine (AZT), Lamivudine (3TC), Ritonavir-boosted Lopinavir (LPV/r), Efavirenz (EFV)

**Table S2.**
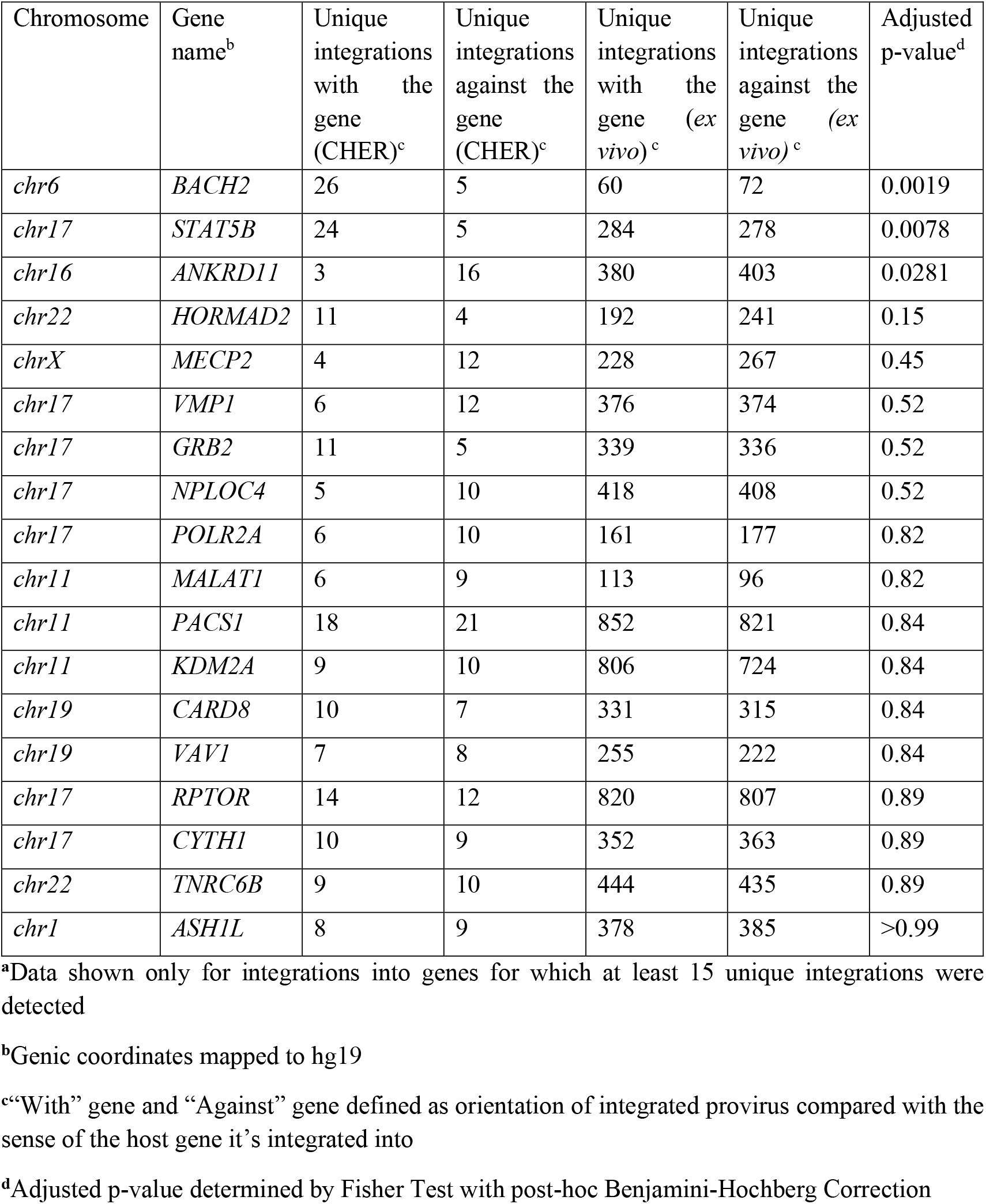
Orientation Bias for Genic Integrations Pre-ART^a^.

| Chromosome | Gene name<sup>b</sup> | Unique integrations with the gene (CHER)<sup>c</sup> | Unique integrations against the gene (CHER)<sup>c</sup> | Unique integrations with the gene (*ex vivo*)<sup>c</sup> | Unique integrations against the gene (*ex vivo*)<sup>c</sup> | Adjusted p-value<sup>d</sup> |
| --- | --- | --- | --- | --- | --- | --- |
| *chr6* | *BACH2* | 26 | 5 | 60 | 72 | 0.0019 |
| *chr17* | *STAT5B* | 24 | 5 | 284 | 278 | 0.0078 |
| *chr16* | *ANKRD11* | 3 | 16 | 380 | 403 | 0.0281 |
| *chr22* | *HORMAD2* | 11 | 4 | 192 | 241 | 0.15 |
| *chrX* | *MECP2* | 4 | 12 | 228 | 267 | 0.45 |
| *chr17* | *VMP1* | 6 | 12 | 376 | 374 | 0.52 |
| *chr17* | *GRB2* | 11 | 5 | 339 | 336 | 0.52 |
| *chr17* | *NPLOC4* | 5 | 10 | 418 | 408 | 0.52 |
| *chr17* | *POLR2A* | 6 | 10 | 161 | 177 | 0.82 |
| *chr11* | *MALAT1* | 6 | 9 | 113 | 96 | 0.82 |
| *chr11* | *PACS1* | 18 | 21 | 852 | 821 | 0.84 |
| *chr11* | *KDM2A* | 9 | 10 | 806 | 724 | 0.84 |
| *chr19* | *CARD8* | 10 | 7 | 331 | 315 | 0.84 |
| *chr19* | *VAV1* | 7 | 8 | 255 | 222 | 0.84 |
| *chr17* | *RPTOR* | 14 | 12 | 820 | 807 | 0.89 |
| *chr17* | *CYTH1* | 10 | 9 | 352 | 363 | 0.89 |
| *chr22* | *TNRC6B* | 9 | 10 | 444 | 435 | 0.89 |
| *chr1* | *ASH1L* | 8 | 9 | 378 | 385 | >0.99 |

<sup>a</sup>Data shown only for integrations into genes for which at least 15 unique integrations were detected
<sup>b</sup>Genic coordinates mapped to hg19
<sup>c</sup>“With” gene and “Against” gene defined as orientation of integrated provirus compared with the sense of the host gene it’s integrated into
<sup>d</sup>Adjusted p-value determined by Fisher Test with post-hoc Benjamini-Hochberg Correction

**Table S3.** Orientation Bias for Genic Integrations On-ART^a^.

| Chromosome | Gene name<sup>b</sup> | Unique integrations with the gene (CHER)<sup>c</sup> | Unique integrations against the gene (CHER)<sup>c</sup> | Unique integrations with the gene (*ex vivo*)<sup>c</sup> | Unique integrations against the gene (*ex vivo*)<sup>c</sup> | p-value<sup>d</sup> |
| --- | --- | --- | --- | --- | --- | --- |
| *chr17* | *STAT5B* | 31 | 6 | 284 | 278 | 6.2E-05 |
| *chr6* | *BACH2* | 12 | 4 | 60 | 72 | 0.034 |

<sup>a</sup>Data shown only for integrations into genes for which at least 15 unique integrations were detected in vivo and at least 1 unique integration ex vivo
<sup>b</sup>Genic coordinates mapped to hg19
<sup>c</sup>“With” gene and “Against” gene defined as orientation of integrated provirus compared with the sense of the host gene it’s integrated into
<sup>d</sup>p-Value determined by Fisher Test – no post-hoc adjustments performed

**Figure S1.**
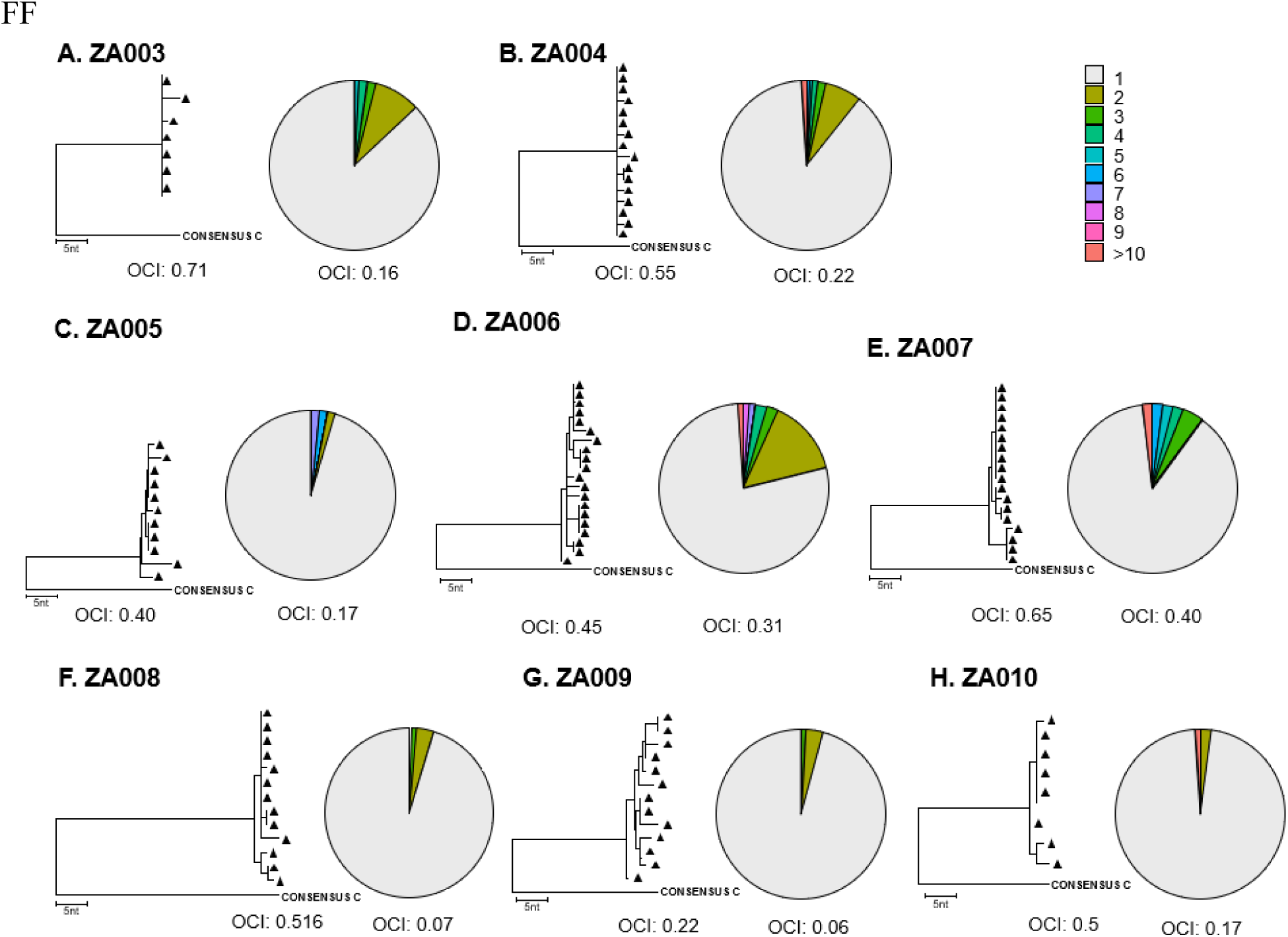
Number of Detections of Integration Sites. For each study participant, a neighbor-joining phylogenetic tree representing gag-pol single genome sequences with its respective OCI value is shown on the left; on the right, a pie chart representing the number of detections of integrations sights by ISA and the respective OCI value.

